# Histone deacetylase-1 is required for epigenome stability in *Neurospora crassa*

**DOI:** 10.1101/2025.01.17.633486

**Authors:** Felicia Ebot-Ojong, Aileen R. Ferraro, Farh Kaddar, Clayton Hull-Crew, Ashley W. Scadden, Andrew D. Klocko, Zachary A. Lewis

**Affiliations:** Department of Microbiology, University of Georgia, Athens, GA, 30602 USA; Department of Chemistry & Biochemistry, University of Colorado Colorado Springs, Colorado Springs, CO, 80918, USA

**Keywords:** constitutive heterochromatin, facultative heterochromatin, Polycomb Repressive Complex 2, H3K27me3, histone deacetylase, histone post-translational modifications

## Abstract

Polycomb group (PcG) proteins form chromatin modifying complexes that stably repress lineage- or context-specific genes in animals, plants, and some fungi. Polycomb Repressive Complex 2 (PRC2) catalyzes trimethylation of lysine 27 on histone H3 (H3K27me3) to assemble repressive chromatin. In the model fungus *Neurospora crassa,* H3K27me3 deposition is controlled by the H3K36 methyltransferase ASH1 and components of constitutive heterochromatin including the H3K9me3-binding protein HETEROCHROMATIN PROTEIN 1 (HP1). Hypoacetylated histones are a defining feature of both constitutive heterochromatin and PcG-repressed chromatin, but how histone deacetylases (HDACs) contribute to normal H3K27me3 and transcriptional repression within PcG-repressed chromatin is poorly understood. We performed a genetic screen to identify HDACs required for repression of PRC2-methylated genes. In the absence of HISTONE DEACETYLASE-1 (HDA-1), PRC2-methylated genes were activated and H3K27me3 was depleted from typical PRC2-targeted regions. At constitutive heterochromatin, HDA-1 deficient cells displayed reduced H3K9me3, hyperacetylation, and aberrant enrichment of H3K27me3 and H3K36me3. CHROMODOMAIN PROTEIN-2 (CDP-2) is required to target HDA-1 to constitutive heterochromatin and was also required for normal H3K27me3 patterns. Patterns of aberrant H3K27me3 were distinct in isogenic Δ*hda-1* strains, suggesting that loss of HDA-1 causes stochastic or progressive epigenome dysfunction. To test this, we constructed a new *Δhda-1* strain and performed a laboratory evolution experiment. Deletion of *hda-1* led to progressive epigenome decay over hundreds of nuclear divisions. Together, our data indicate that HDA-1 is a critical regulator of epigenome stability in *N. crassa*.

## Introduction

Formation of specialized heterochromatin is directed by histone post-translational modifications (PTMs) (1). In some cases, repressive chromatin states can be epigenetically inherited over the lifetime of an organism. Different types of repressive chromatin domains are enriched for characteristic histone modifications and have distinct functions. Constitutive heterochromatin (cHet) is found at repeat-rich regions of the genome and is typically enriched for di- or tri-methylated lysine-9 on histone H3 (H3K9me2/3) (1). A second type of repressed chromatin is assembled and maintained by Polycomb Group (PcG) proteins, which interact to form Polycomb Repressive Complex (PRC) 1 and PRC2 (2–4). PRC2 methylates histone H3 lysine 27 in PcG-repressed chromatin, which is often referred to as facultative heterochromatin (fHet) because PcG-repressed genes are expressed in context- or cell-type-specific manner (5–7). Notably, hypoacetylated histones are a characteristic feature of both cHet and fHet domains (1, 8).

Defective assembly or maintenance of heterochromatin is associated with human diseases including cancer, highlighting the critical need to define genes and mechanisms that structure the epigenome (9, 10). The PcG proteins have been intensely studied in diverse experimental systems, revealing that numerous interconnected regulatory mechanisms establish and maintain H3K27me3-enriched domains. In plants and mammals, accessory proteins can recruit PRC1 or PRC2 to target sites (2, 3, 11–15). In addition, the activity of PRC2 is modulated by chromatin features, including activating chromatin modifications (16–18), inhibitory modifications (19–21), RNA (22–25), and densely packed chromatin (26).

The fungus *Neurospora crassa* encodes conserved components of both cHet and fHet pathways and is a well-established model system to investigate assembly and maintenance of heterochromatin (5, 27). Prior studies with *N. crassa* showed that normal H3K27 methylation depends on core PRC2 components (28) and a variety of additional factors including H2A.Z (29), accessory proteins (CAC-3/NPF, EPR-1, and PAS) (28, 30, 31), the chromatin remodeler IMITATION SWITCH (ISWI) (32, 33), and other histone modifications such as H3K36me3 (34), and H3K9me3 (35, 36). In contrast to mammalian systems, H3K36 methylation frequently co-occurs with and can promote H3K27me3 in fungi (37, 34, 38, 39). Thus, PRC2 localization and activity is influenced by diverse histone modifications in *N. crassa*, like in higher eukaryotes, but much remains unknown about how various modifications cooperate to structure the epigenome.

In plants, animals, and fungi, molecular features of cHet antagonize H3K27me3 (40). In wild type *N. crassa,* cHet domains are defined as repeat-rich, gene-poor DNA sequences enriched for H3K9me3, HP1, DNA cytosine methylation (5mC), and hypoacetylated histones (41, 27, 42). cHet domains depend on multiple HP1-containing complexes that bind to H3K9me3 and modulate cHet structure by directing histone deacetylation, 5mC deposition, or by recruiting the anti-silencing factor DNA Methylation Modulator 1 (DMM-1) (43–47, 42, 48). In *N. crassa* mutants lacking H3K9me3 or HP1, H3K27me2/3 is lost from native PRC2 target domains and aberrantly gained at cHet regions (35, 36). Similar redistribution of H3K27me3 is observed in mammalian cells depleted for H3K9me3 or 5mC (49–52). In mutant mouse embryonic stem cells (mESCs) lacking 5mC, a non-canonical PRC1 complex (ncPRC1.1) binds to hypomethylated CpG-rich DNA in pericentromeric cHet regions and deposits ubiquitylated H2A (H2Aub), leading to PRC2 recruitment and H3K27 methylation of H3K9me3-depleted histones (53, 54). Thus, features of mammalian cHet domains modulate H3K27me3 localization in at least two ways; 5mC prevents aberrant recruitment of PRC1 and PRC2 to mammalian cHet domains, whereas H3K9me3 inhibits PRC2 activity at these regions. In contrast to mammals, loss of 5mC does not impact H3K27me2/3 patterns in *N. crassa* (35, 36), and how H3K9me3 and HP1 exclude H3K27me3 from cHet regions is not understood. In addition, it is unclear if other components of cHet domains, such as histone deacetylases, impact H3K27me3 localization in any system.

Here, we show that HDA-1, a cHet-specific histone deacetylase, is required for repression of PRC2-methylated genes in *N. crassa.* HDA-1 is a component of the <u>H</u>P1/<u>C</u>DP-2/<u>H</u>DA-1/<u>C</u>HAP (HCHC) complex, which limits DNA accessibility at cHet domains and is critical for normal three-dimensional (3D) chromosome conformation (42, 48, 55, 56). In cells lacking HCHC components, we found that H3K27me3 is lost from typical PRC2-methylated genes (fHet) and gained at cHet regions. We observed distinct patterns of H3K27me3 in strains lacking individual HCHC components and in individual Δ*hda-1* strain isolates, raising the possibility that loss of HDA-1 targeting or activity leads to stochastic or progressive changes to the epigenome. To test this hypothesis, we constructed a new Δ*hda-1* strain and performed a lab evolution experiment. In this new Δ*hda-1* strain, changes in the pattern of H3K27me3 were progressive, taking place over hundreds of nuclear divisions, and were accompanied by altered lysine acetylation, H3K36me3, and H3K9me3. Together, our results show that HCHC is a critical regulator of epigenome stability in *N. crassa*.

## Materials and Methods

### Strains, strain construction, and growth conditions

Strains and primers are listed in Tables S1 and S2, respectively. Strains were grown in Vogel’s minimal medium (VMM) with 1.5% sucrose or with 0.05% fructose, 0.05% glucose, and 2% sorbose as described (57). Media was supplemented with 25 mg/mL histidine or 200µg/mL hygromycin B where appropriate. Gene deletion strains were obtained from the *Neurospora* knockout collection (58), unless otherwise indicated. The NKL9 and NKL10 Δ*hda-1* strains were generated in the Klocko lab by backcrossing an Δ*hda-1* strain (FGSC 12003) to wild type and passaging the resulting progeny. A new *hda-1* knockout mutant, strain S863, was constructed for this study using a split marker approach (59). Partial deletion fragments were generated by polymerase chain reaction (PCR) using genomic DNA from an existing Δ*hda-1* strain (S297) as a template using primer pairs *hda*-1 5’ flank FP + *hph* SM-1 and *hda-1* 3’ flank RP + *hph* SM-2. Fragments were transformed into the wild type strain S2 by electroporation as described (60). To perform complementation experiments, we first generated plasmid pBM61_HDA-1-3xFLAG by cloning synthetic gene fragments (Twist Biosciences) corresponding to the wild type *hda-1-3xflag* sequence under control of the native *hda-1* promoter (546bp upstream of the annotated transcription start site) into the plasmid backbone of pBM61-3xFLAG amplified by PCR using primers pBM61_Inverse P3 + pBM61_Inverse P2. A plasmid containing a presumptive catalytic dead *hda-1^h225A^-3xflag* allele was constructed in the same way. Plasmid sequences were verified by sequencing (Plasmidsaurus). Both constructs were introduced into the *his-3* locus of strain S188 (*his-3;* Δ*hda-1*) as described (60).

### Chromatin Immunoprecipitation (ChIP), CUT&RUN, RNA-seq, and Hi-C

All antibodies used in this study are reported in Table S3. ChIP-seq was performed as described (56, 61, 62). For RNA-seq experiments, RNA isolation and RNA-seq library preparation was performed as described (Kamei et al., 2021; Zhou et al., 2018). We adapted CUT&RUN for *N. crassa* by modifying a previously published protocol (64). For Neurospora CUT&RUN, cells were grown in 5mL VMM + 1.5% sucrose for 16-18 hours. Mycelial mats were harvested using a Buchner funnel and washed with 10mL of PBS. Washed mycelial mats were transferred to a clean petri dish, flooded with 1mL of nuclear isolation buffer (20 mM HEPES pH 7.5, 0.5 mM Spermine tetrahydrochloride, 80 mM KCl, 20 mM NaCl, 0.1 mM CaCl_2_, 15 mM 2-Mercaptoethanol, 0.3% Triton X-100), and chopped with a razor blade for 2 minutes. The suspension of mycelial fragments was filtered through Miracloth (Millipore-Sigma, Catalog # 475855) into a 1.5 mL microfuge tube and centrifuged for 5 minutes at 1000 x G to remove cell debris. The nuclei-containing supernatant was subject to CUT&RUN using an established procedure (65).

*In situ* Hi-C was performed using crosslinked nuclei. Briefly, 25mL cultures of 1x Vogel’s and 1.5% sucrose were started from ∼1.0 x 10^8^ conidia and grown overnight at 32°C with shaking at 200 rpm. Cultures were crosslinked with 1% [final] formaldehyde, quenched with 125mM Tris-HCl, pH 8.0, and the mycelial mat was isolated by vacuum filtration and placed in a petri dish on ice. Nuclear isolation buffer (5mL; 20mM HEPES pH 7.5, 80mM KCl, 20mM NaCl, 0.1mM CaCl_2_, 15mM β-mercaptoethanol, 0.3% Triton, and 0.5mM Spermine tetrahydrochloride [freshly added]) was added and the mycelial mat was rapidly cut with a new razor blade. The lysate was filtered through autoclaved Miracloth (Millipore Sigma, cat# 475855-1R) lining a funnel into a 15mL conical tube, and nuclei were collected by centrifugation at 4,500 rpm for 5 minutes. The pellet was washed with an additional 1mL nuclear isolation buffer, and nuclei pellets were stored at −80°C. Nuclei comprising at least 3.5 ug of DNA were then used for the remainder of the *in situ* Hi-C protocol using the restriction enzyme *Mse*I to more efficiently capture AT-rich genomic loci, starting with 6.25% SDS treatment to make the nuclei porous, as previously described (56). All DNA sequencing data was generated as paired-end reads on a NextSeq 2000, NovaSeq 6000, or a NovoSeq X instrument.

### Analysis of High-throughput sequencing data

For ChIP-seq, fastq files were trimmed to remove adaptor sequences using TrimGalore! V0.6.(66), and then mapped to *N.crassa* assembly 12 (GCA_000182925.2) or assembly 14 for comparison to Hi-C data (67) with BWA v0.7.17 mem (option -M) (68) or bowtie2 (69), respectively. Reads were sorted and indexed using SAMtools v1.16.1 (70). Bigwig files for visualization were made using deepTools v3.5.2 ‘bamCoverage’ module (71), normalized by either BPM (bins per million mapped reads) or RPKM (reads per kilobase per million mapped reads), while heatmaps of ChIP-seq data were created using deepTools ‘computeMatrix’ and ‘plotHeatmap’ modules. Resulting bigwig files were visualized with the Integrative Genome Viewer (72), and saved images were used for figure creation. The ‘csaw’ and ‘Complex Heatmap’ packages in R were used to compute and plot H3K27me3, H3K9me3, H3K36me3, or lysine acetylation (Kac) ChIP-seq enrichment values in wild type and passaged Δ*hda-1* strains (73). Previously published ChIP-seq datasets used for this manuscript include wild type H3K9me3 (NCBI GEO submission # GSE232933) and wild type H3K27me2/3 (merged files from GSE68897 and GSE100770) (36, 56, 74).

For RNA-seq, reads were trimmed with TrimGalore v0.6.5 (66) and mapped using STAR v2.7.10 (75). Output files in BAM format were indexed using SAMtools v1.9, and alignment counts were determined using subRead v2.0.1 ‘featureCounts’ (76). Differential expression was calculated using DESeq2 (77). PRC2-targeted genes defined previously and are listed in (**Dataset S1**) (32).

For Hi-C data processing, output fastq datasets were mapped to the *N. crassa* reference genome, version 14 (nc14) (67) with Bowtie2 (69), and contact matrices were built at a high (1 kb) resolution using the “hicBuildMatrix” command of the hiCExplorer software package; lower resolution (10 kb or 20 kb bin) matrices were generated with the hicMergeMatrixBins command. (78). To compare to the previously published wild type *Mse*I Hi-C dataset, a number of total reads were extracted from the NKL9 *Mse*I dataset that would give approximately equal numbers of valid reads for comparison to a wild type N150 *Mse*I Hi-C dataset (or reads were extracted from the wild type *Mse*I dataset when comparing to NKL10) using the sed command (67). After mapping reads and building the contact matrix with the selected fastq files, the resulting contact matrices were compared using the hicCompareMatrices command of hiCExplorer. Images were generated with the hicPlotMatrix command from hiCExplorer and used for figure construction.

## Results

### HDA-1 is required for repression of PRC2-targeted genes

To determine if any histone deacetylases are required for transcriptional repression of genes in facultative heterochromatin (fHet; i.e. H3K27me3-enriched domains), we tested available HDAC-deficient mutants for defective gene silencing. We performed RNA-seq to quantify mRNA levels in wild type and ten gene deletion strains lacking predicted histone deacetylase enzymes (79). The *N. crassa hda-3* gene is essential and was not included in this study (80). For all HDAC-deficient strains tested, reproducible results were obtained with replicate RNA-seq samples (**Figure S1**). For each gene deletion strain, we first examined average expression levels of previously defined PRC2-methylated genes (32), which are tightly repressed in vegetative mycelia (Deaven et al. in preparation; **Supplemental Dataset 1**). The Δ*hda-1* strain showed striking upregulation of PRC2-targeted genes relative to wild type or other HDAC-deficient mutants (**Figure 1A**). The extent of upregulation in Δ*hda-1* was similar to that observed for the Δ*set-7* control strain, which lacks all H3K27me3. A heatmap plotting relative expression of each PRC2-targeted gene confirmed broad upregulation of this gene set in Δ*hda-1* but not in other HDAC-deficient mutants (**Figure 1B**). In total, 2263 genes had significant expression changes in the Δ*hda-1* mutant compared to wild type (**Figure 1C, Dataset S2;** +/-2-fold [Δ*hda-1/*WT], adjusted p-value of < 0.05). Of these, 1493 genes displayed increased expression whereas 770 genes showed reduced expression levels compared to wild type. More than 30% of PRC2 target genes were significantly upregulated in the Δ*hda-1* by more than 2-fold (**Dataset S2**). Importantly, genes encoding PRC2 complex members or other known components of the fHet pathway did not exhibit altered expression in the Δ*hda-1* strain (**Dataset S2**). We conclude that HDA-1 is required for normal repression of ∼15% of *N. crassa* genes, including a significant fraction of silent PRC2-methylated genes located in fHet regions.

**Figure 1:**
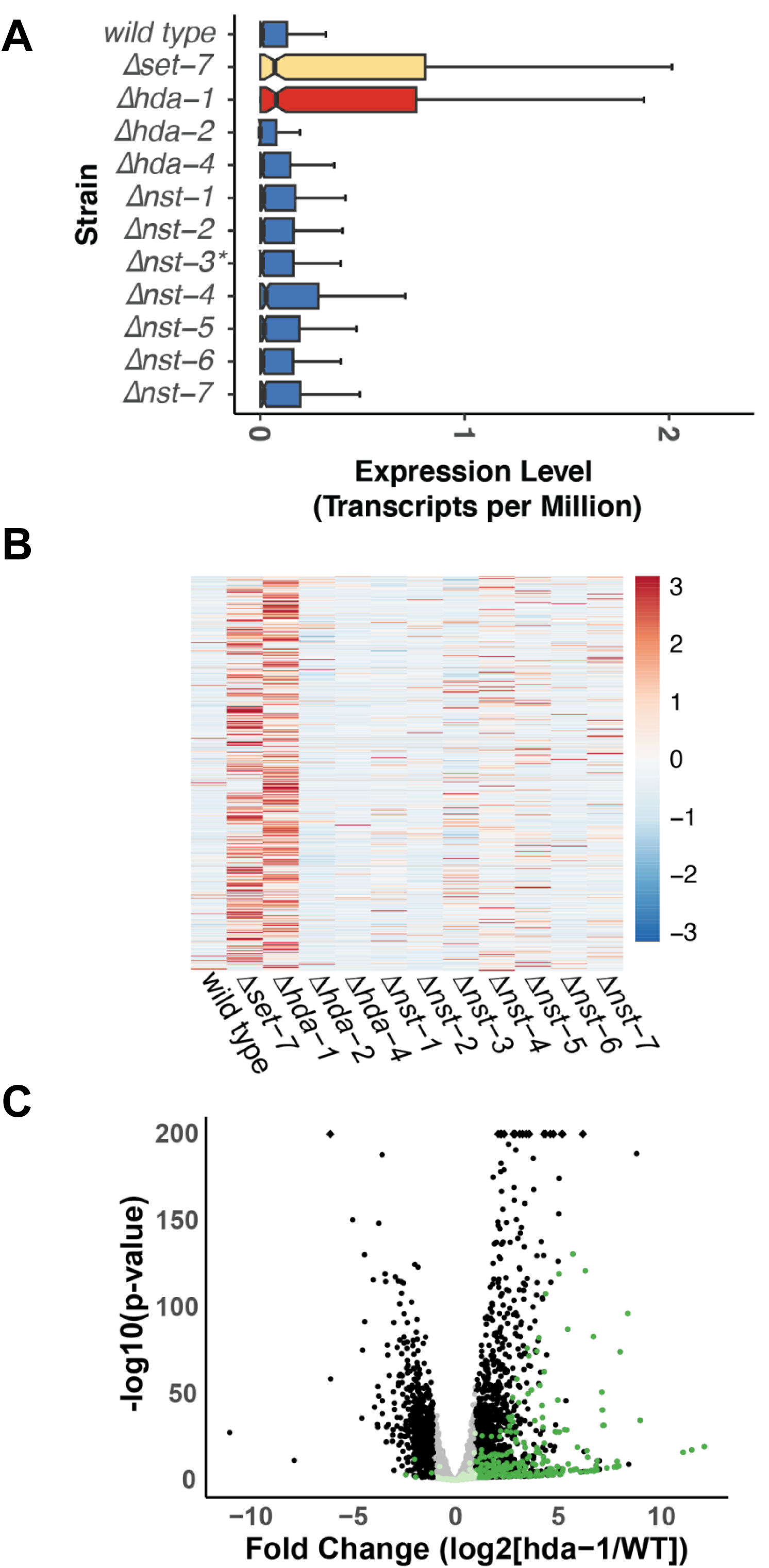
HDA-1 is required for the repression of PRC2-methylated genes. (**A**) The boxplots show the average level of expression for all PRC2-target genes (n= 679) for the indicated strains, with the x-axis values corresponding to log2-transformed transcripts per million (TPM) read counts. Lines represents the median expression value, and the notches indicate the 95% confidence interval. (**B**) The heatmap shows relative expression levels of each PRC2-target gene for the indicated strains (n= 537 genes expressed in at least one strain). (**C**) The volcano plot shows differentially expressed genes for wild type and Δ*hda-1*. Dark green spots correspond to PRC2-methylated genes that meet fold-change and statistical thresholds (log2[fold change] +/-1.5 and an adjusted p-value < 0.05). Black spots correspond to non-PRC2 methylated genes that meet the same fold-change and statistical threshold. Genes that do not meet statistical or fold-change thresholds are colored in pale green for PRC2-target genes or grey for non-PRC2 targets.

### Histone modification patterns are altered in *the* ***Δ***hda-1 strain

To determine if HDA-1 is required for normal chromatin structure at fHet domains, we performed CUT&RUN to examine the distribution of H3K27me3 across the *N. crassa* genome. Because HDA-1 is reportedly targeted to cHet regions by HP1 (42), we also examined the distribution of H3K9me3. **Figure 2A** shows the distributions of H3K27me3 and H3K9me3 across Linkage Group III for wild type and Δ*hda-1*. We observed substantial reduction of H3K27me3 in fHet regions (i.e. domains enriched for H3K27me3 in a wild type strain). Loss of H3K27me3 at fHet domains was accompanied by gains of H3K27me3 in cHet regions (i.e. domains enriched for H3K9me3 in a wild type strain). For cHet domains that gained H3K27me3, we observed lower levels of H3K9me3 enrichment compared to cHet regions that did not gain H3K27me3 (**Figure 2A-B**). Replicate experiments produced similar results (**Figure S2A**). To examine the global changes in the repressive chromatin landscape, we generated heatmaps to visualize H3K27me3 and H3K9me3 levels across all wild type fHet and cHet domains (**Figure 2C-D**). This confirmed widespread loss of H3K27me3 from fHet domains, while ∼ 20% of cHet regions gained H3K27me3 and lost H3K9me3.

**Figure 2:**
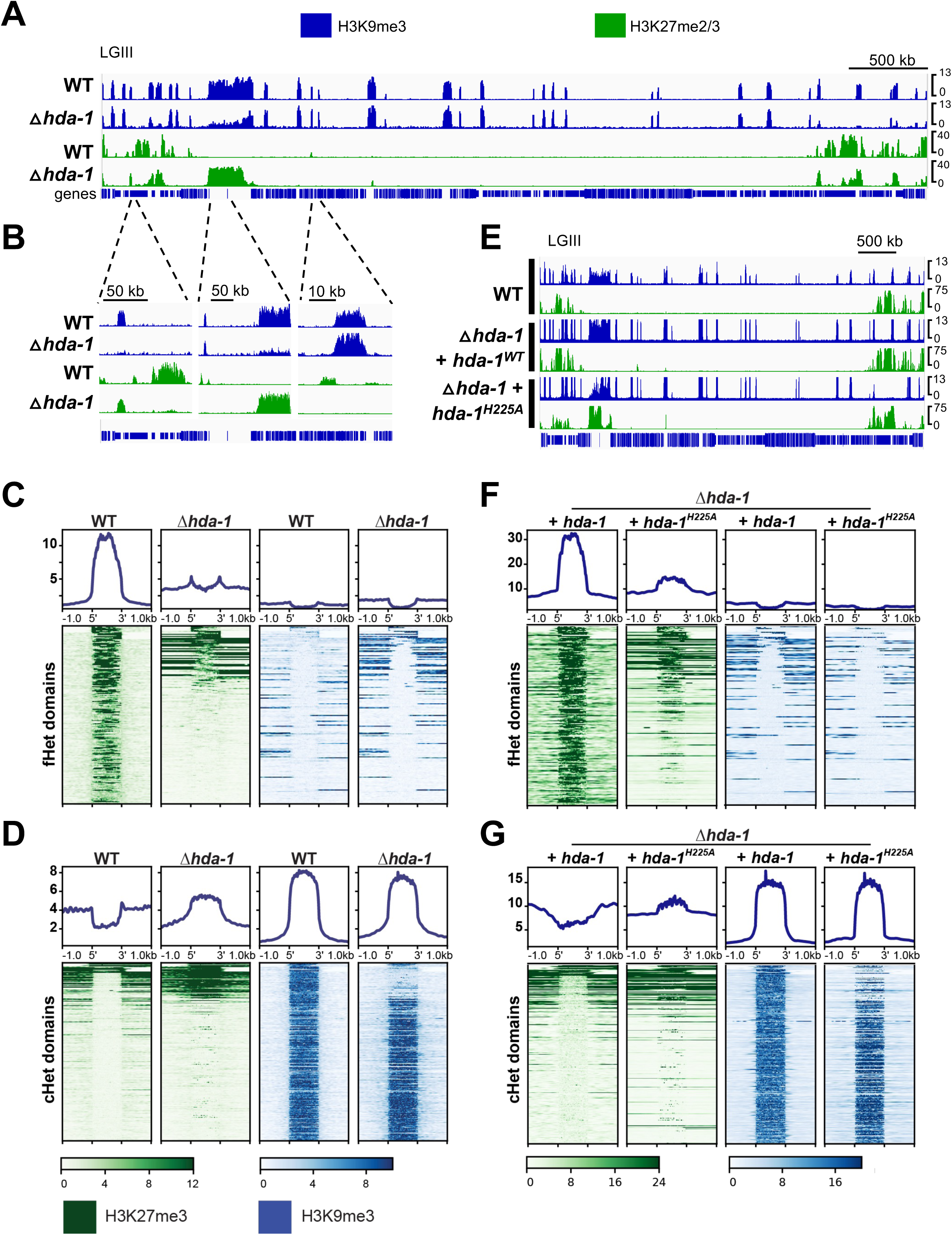
H3K27me3 aberrantly localizes to heterochromatin in the Δ*hda-1* mutant. (**A-B**) The genome browser image shows the distribution of H3K9me3 (blue) and H3K27me3 (green), as assessed by CUT&RUN, across (**A**) Linkage Group III or (**B**) three representative regions shown at higher magnification. The scale bars indicate genomic distance in kilobases (kb). (**C-D**) Heatmaps depicting H3K27me3 (green) or H3K9me3 (blue) enrichment across (**C**) all wild-type fHet domains or (**D**) all wild-type cHet domains for the indicated strains. The heatmaps are sorted such that the first 14 rows in each heatmap correspond to the 14 subtelomeric regions located at the chromosome ends. The remaining rows are sorted by the level H3K27me3 enrichment in Δ*hda-1* in descending order from highest to lowest. Each peak size is normalized to 100% with 1,000 bp shown up/downstream. **(E**) The genome browser tracks show the distribution of H3K9me3 (blue) and H3K27me3 (green), as in A, for the indicated strains. (**F-G**) The heatmaps show enrichment of H3K27me3 (green) or H3K9me3 (blue) across (**F**) all wild-type fHet domains or (**G**) all wild type cHet domains for the indicated strains, ordered as in panels C-D.

To confirm that changes in histone PTMs are caused by loss of HDA-1, we introduced an *hda-1-3xflag* construct into an Δ*hda-1* deletion strain and performed ChIP-seq. Wild-type patterns of H3K27me3 and H3K9me3 were restored in this complemented strain (**Figure 2E-F**). Importantly, ChIP-seq data validated results obtained using CUT&RUN (**Figure 2A-D)**, which had not previously been adapted for *N. crassa.* We next asked if HDA-1 catalytic activity is required for normal patterns of H3K27me3 and H3K9me3. We generated a mutant *hda-1^H225A^-3xflag* construct in which a histidine required for catalytic activity is mutated to alanine (81) and introduced this allele into Δ*hda-1*. This mutant *hda-1* allele failed to rescue H3K9me3 or H3K27me3 patterning defects (**Figure 2E,G**). Both epitope-tagged proteins were expressed at similar levels (**Figure S2B**), and replicate ChIP-seq experiments produced similar results (**Figure S2C**). We conclude that *hda*-1 is required for normal patterns of both H3K27me3 and H3K9me3 in *N. crassa*.

HDA-1-deficient cells exhibit hyperacetylation at constitutive heterochromatin domains (42, 48, 56). To explore the relationship between lysine acetylation and altered fHet/cHet histone PTMs in the Δ*hda-1* strain, we performed ChIP-seq using antibodies to pan acetyl-lysine (Kac), H3K27me3, and H3K9me3. We used a pan-acetyl lysine antibody because previous work revealed a striking increase in H2B acetylation in Δ*hda-1* strain (82), but no H2Bac antibodies are available for *N. crassa.* In WT, low acetylation levels were observed in cHet domains marked by H3K9me3 and to a lesser extent, in fHet domains marked by H3K27me3 (**Figures 3A; S3A**). These results are similar to those obtained in prior studies using antibodies to acetylated H3 residues (42, 56, 80). In the Δ*hda-1* strain, we observed hyperacetylation in cHet regions, such as the centromere, consistent with previous reports of the HCHC targeting these loci (**Figure 3A, S3A**) (42).

**Figure 3:**
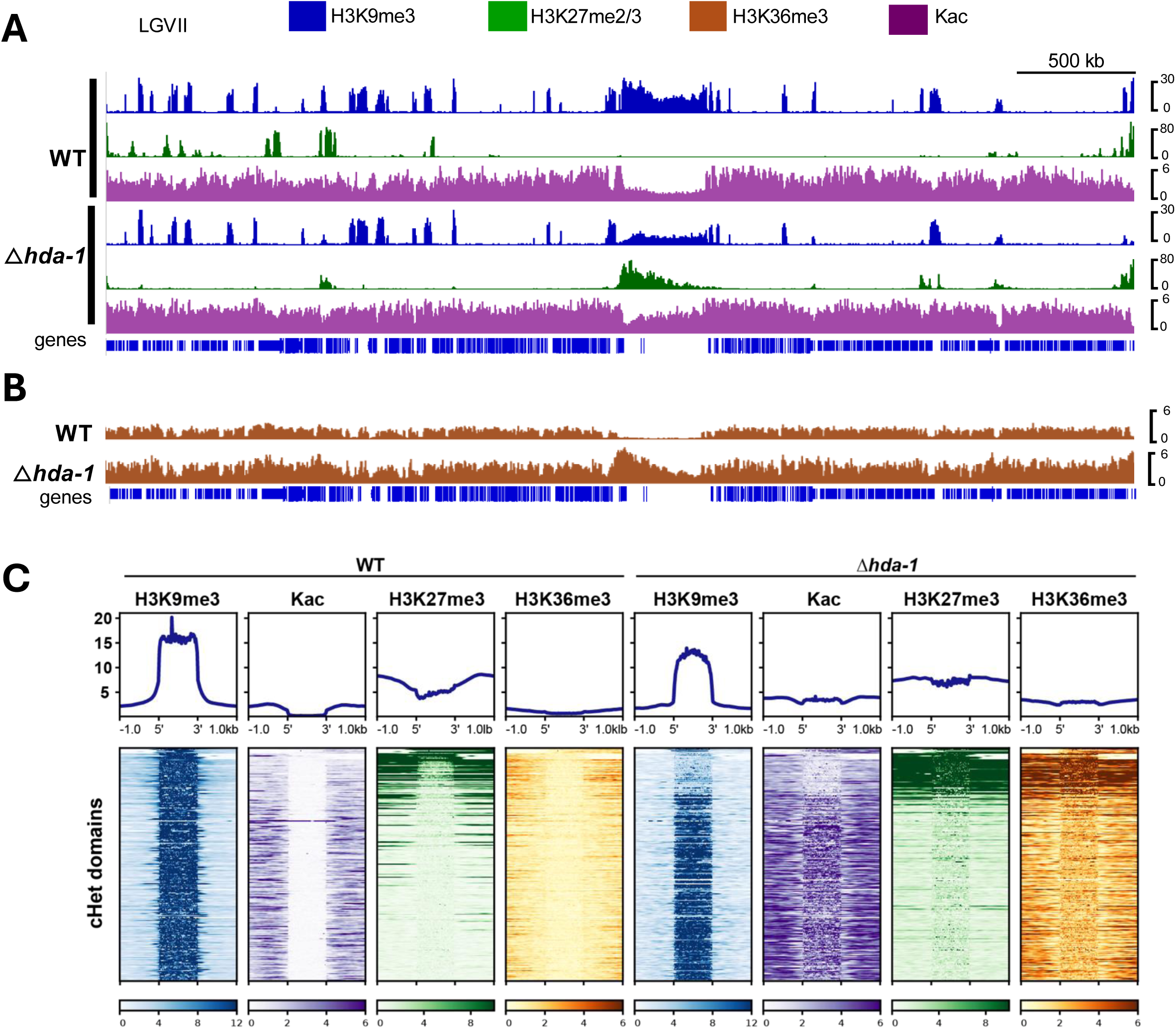
Acetylation and H3K36me3 patterns are altered in Δ*hda-1*. (**A-B**) The genome browser tracks show ChIP-seq enrichment of (**A**) H3K9me3 (blue), H3K27me3 (green), and pan-acetyl lysine (Kac; purple) or (**B**) H3K36me3 across Linkage Group VII, as in Figure 2A. (**C**) The heatmaps show ChIP-seq enrichment of H3K9me3 (blue), H3K27me3 (green), H3K36me3 (orange), or pan-acetyl lysine (H3Kac; purple) for wild type (left) or Δ*hda-1* (right) across all cHet domains. The heatmaps are scaled and sorted as in Figure 2C-D, with subtelomeres at the top followed by the descending H3K27me3 signal in fHet regions in a Δ*hda-1* strain.

In *N. crassa*, H3K27me3 co-occurs with ASH1-dependent H3K36 methylation. In fHet regions, ASH-1 deposits H3K36me2, which is efficiently converted to H3K36me3 by SET-2 (34, 83). SET-2 also methylates H3K36me1/2/3 in the coding sequences of actively transcribed genes within euchromatin. We performed ChIP-seq to determine if HDA-1 is required for normal H3K36me3 localization (**Figure 3B**). Consistent with previous results, wild type H3K36me3 patterns were distinct in euchromatin, fHet, and cHet regions. In euchromatin, H3K36me3 was restricted to gene bodies. In fHet domains, H3K36me3 overlapped with H3K27me3 covering both promoters and gene bodies across large, multi-gene regions. In cHet regions, no H3K36me3 enrichment was observed in wild type. In the Δ*hda-1* strain, H3K36me3 was enriched in a subset of cHet domains, similar to the aberrant pattern of H3K27me3 in this mutant. We examined the interrelationships of Kac, H3K27me3, H3K36me3, and H3K9me3 by plotting enrichment of these PTMs across all wild type cHet domains (**Figure 3C**). As expected, wild-type cHet regions were heavily enriched for H3K9me3 and devoid of Kac, H3K36me3, and H3K27me3, except for small domains adjacent to each telomere that exhibit co-enrichment of fHet- and cHet-specific PTMs (84). In Δ*hda-1*, individual cHet domains gained either Kac or both H3K27me3 and H3K36me3. The subset of cHet domains that gained H3K27me3 and H3K36me3 exhibited concomitant loss of H3K9me3 (∼20% of cHet regions). In contrast, most cHet domains (∼80%) displayed typical enrichment of H3K9me3 accompanied by aberrant hyperacetylation. The patterns of Kac and H3K36me3 in euchromatin were unaffected in the Δ*hda-1* mutant, consistent with the known role of HDA-1 at cHet (**Figure S3C-D**). Replicate ChIP-seq samples produced similar results (**Figure S3E**). Thus, loss of HDA-1 leads to aberrant deposition of two modifications typically found at fHet regions, H3K27me3 and H3K36me3, in a subset of cHet loci. In addition, reduced acetylation at these cHet loci suggests these fHet modifications can direct recruit a compensatory deacetylase to cHet loci of Δ*hda-1* strains.

### HCHC-deficient mutants exhibit distinct patterns of aberrant H3K27me3

HDA-1 is a component of the HP1-containing HCHC complex. HP1 was previously shown to exclude H3K27me3 from cHet regions (35, 36). We next asked if other components of HCHC are required for normal H3K27me3 localization. We performed CUT&RUN to map enrichment of H3K27me2/3 in strains lacking the individual HCHC components HP1, CDP2, or CHAP (**Figure 4A**). HP1-deficient (Δ*hpo)* and CDP-2-deficient (Δ*cdp-2)* strains exhibited widespread re-localization of H3K27me2/3 from typical PRC2-target domains to cHet domains, while the CHAP-deficient (Δ*chap*) strain had a wild type H3K27me2/3 pattern. Interestingly, the global relocalization pattern of H3K27me2/3 was distinct in the Δ*hpo,* Δ*hda-1,* and Δ*cdp-2* strains. This was evident when H3K27me2/3 enrichment was plotted across all seven *N. crassa* centromeres (**Figure 4A**-**B**). We also plotted H3K27me2/3 enrichment across all wild type cHet domains in each of the HCHC-deficient mutants, which confirmed that the pattern of aberrant H3K27me2/3 enrichment was distinct in each of these strains (**Figure 4C**). Replicate experiments produced similar results (**Figure S4**).

**Figure 4:**
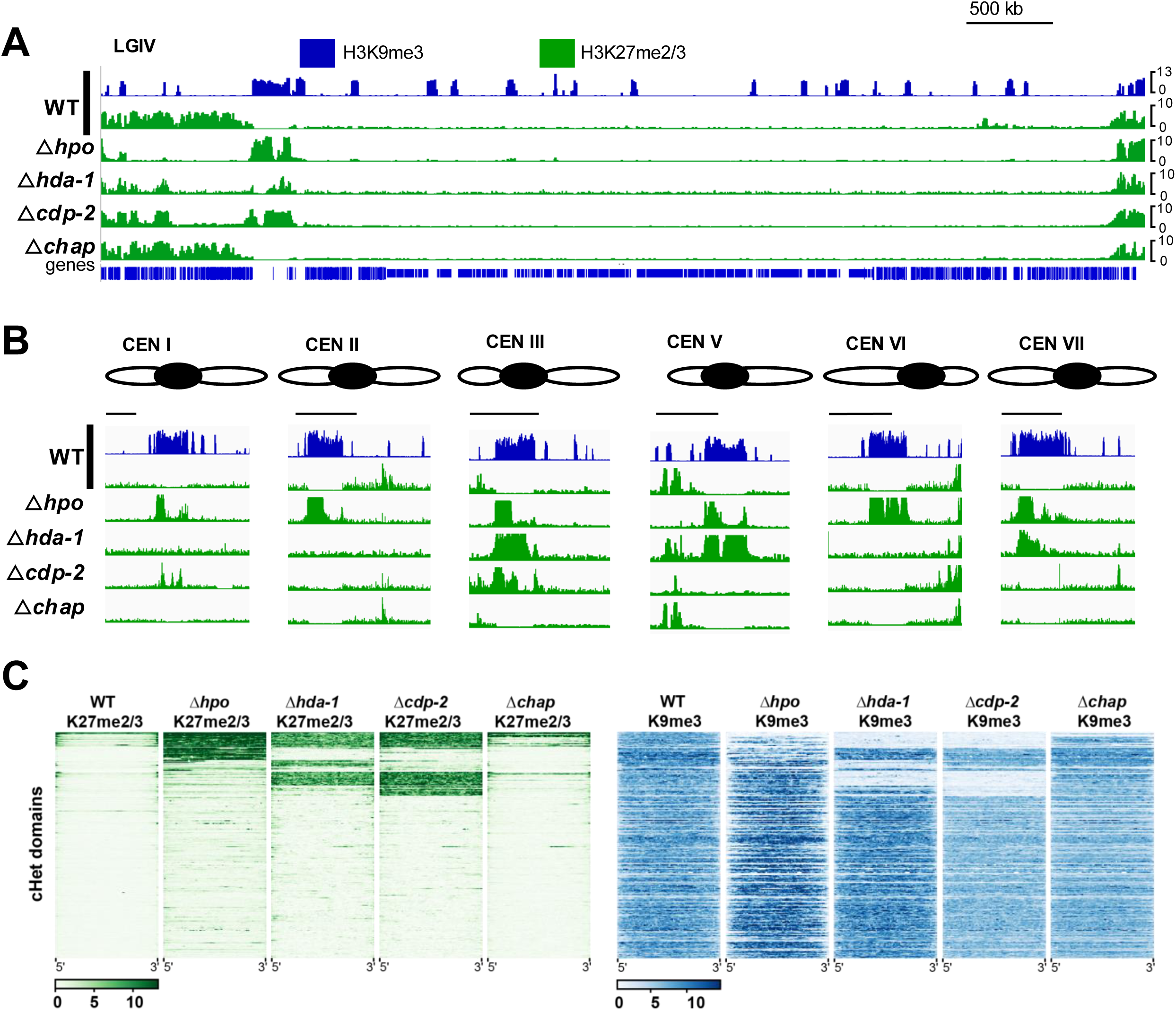
HP1 and CDP-2, but not CHAP, are required for normal H3K27me3 distribution. **(A-B)** The genome browser tracks show CUT&RUN enrichment of H3K9me3 (blue) and H3K27me2/3 (green) across (**A**) Linkage Group IV, or (**B**) the centromeres of the remaining chromosomes, for the indicated strains, as in Figure 2A-B. (**C**) The heatmaps show H3K27me2/3 (green) or H3K9me3 (blue) CUT&RUN enrichment for the indicated strains across all wild type cHet regions, except for 14 sub-telomeric domains adjacent to the chromosome ends. Each cHet region is normalized to 100% size, and no flanking sequences are plotted.

### ***Δ***hda-1 strains have increased intra- and inter-heterochromatin interactions in the nucleus

To assess if altered enrichment of histone PTMs in Δ*hda-1* influences the 3D organization of the *Neurospora* genome, we performed cHet-specific chromosome conformation capture coupled with high-throughput sequencing (*in situ* Hi-C); here, the use of the restriction enzyme *Mse*I (recognition sequence: T^TAA) effectively captures contacts of the AT-rich cHet regions (67, 85). The *Neurospora* genome forms a Rabl chromosome conformation characterized by a centromere cluster and multiple telomere bundles (85–87). The other ∼200 cHet regions strongly associate over Megabases (Mb) of genomic distance. Previously, it was shown that intra- and inter-heterochromatic region interactions are strongly increased in a Δ*cdp-2* strain due to histone hyperacetylation (56). To determine if similar changes are observed in another HCHC-deficient mutant, we performed *in situ* Hi-C using *Mse*I in two independent strain isolates of Δ*hda-1* (NKL9, NKL10). We also performed ChIP-seq for H3K9me3 and H3K27me2/3 in both strains.

As previously shown, *Mse*I Hi-C of a wild type strain produced 20 kilobase (kb) resolution contact probability heatmaps in which the interspersed cHet regions show strong intra-region (on-diagonal) interactions (**Figure 5A**) (67). Moreover, long-range interactions between cHet regions are readily observed, as shown for two individual chromosomes (**Figure 5A,E**; Linkage Groups [LGs] II and V). Similar contact probability heatmaps are observed for the NKL9 Δ*hda-1* strain, except that strong inter-region interactions between cHet domains are more apparent (**Figure 5B,F**), especially for the centromeres of each LG, which are more apt to interact with distal cHet regions. Comparison of the wild type contact probability heatmap to that of the Δ*hda-1* strains showed increased interactions at the center of each cHet region, as well as between cHet regions, with the contacts between centromeres and interspersed heterochromatic regions having the most prominent gains (**Figure 5C,G**). Similar results were previously reported for a Δ*cdp-2* strain (56). These increased cHet domain contacts of the LG II and LGV centromeres in the Δ*hda-1* strain NKL9, relative to a wild type strain, are readily apparent even at a higher resolution (10 kb bins; **Figure 5D,H**). A replicate Hi-C experiment with a second Δ*hda-1* strain isolate, NKL10, showed similar cHet contact probability gains (**Figure S5**). These data suggest that loss of HCHC leads to promiscuous interactions between distant cHet regions, presumably due to hyperacetylation at these regions.

**Figure 5:**
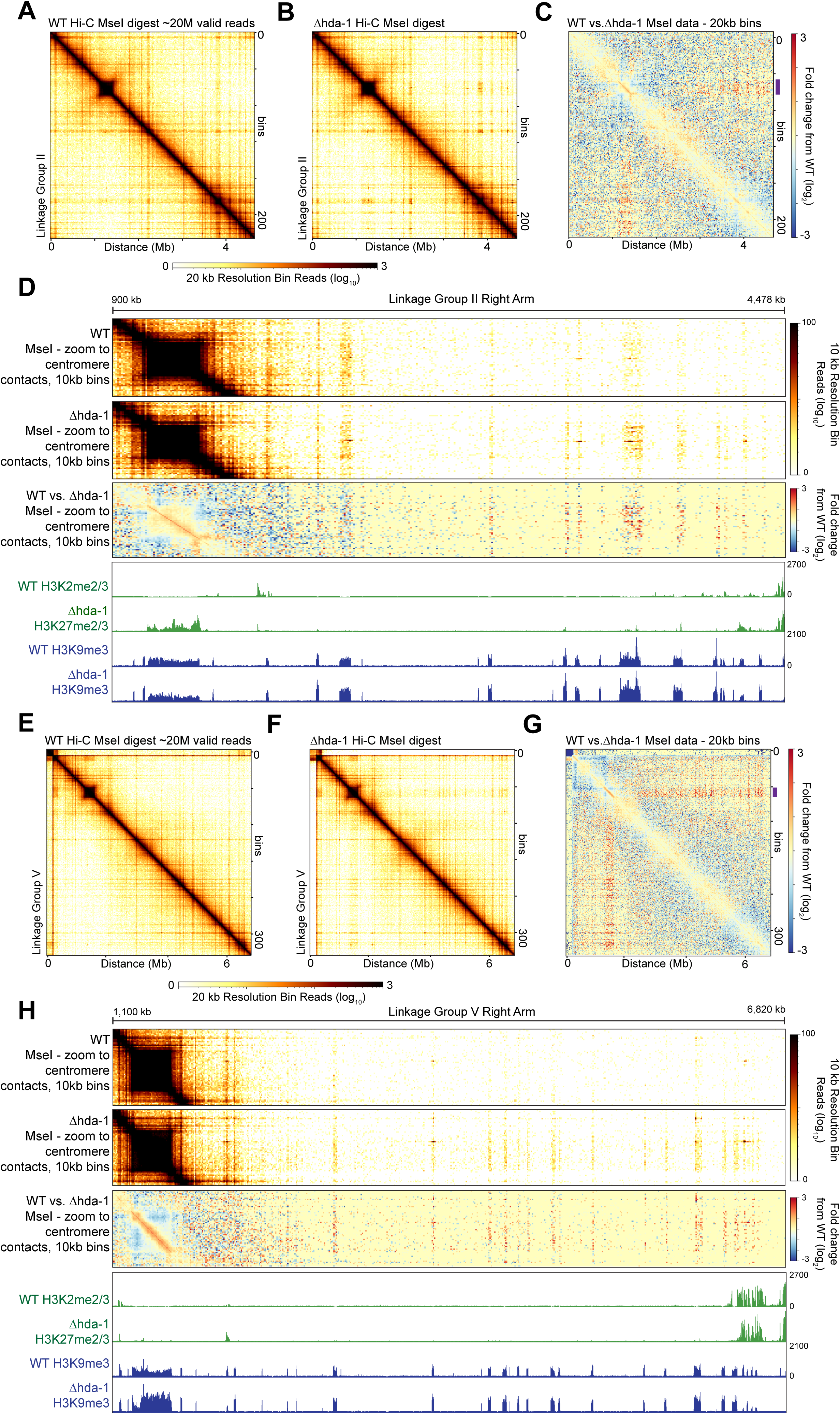
Inter-heterochromatic region contact gains and variable H3K27me2/3 enrichment occur in a Δ*hda-1* strain. Heterochromatin-specific (*Mse*I) *in situ* Hi-C and H3K27me2/3 or H3K9me3 ChIP-seq of wild type or Δ*hda-1* (NKL9) strains are displayed across two Linkage Groups (LG), LG II (**A-D**) or LG V (**E-H**). Contact probability heatmaps of raw *in situ* Hi-C reads at 20 kb resolution of a wild-type strain (**A, E**) or the NKL9 Δ*hda-1* strain (**B, F**) are shown; the heatmap scalebar is below, while the genomic distance is indicated on the x-axis and the number of bins indicated on the right. **(C, G**) Heatmaps showing the change in contact probability (log_2_) between wild type and Δ*hda-1* strains, with the heatmap scalebar on the right. The purple line shows the centromeric region highlighted in panels D and H. (**D, H**) Heatmaps of the contact probability of wild type (top) or Δ*hda-1* (NKL9) strains (middle), or the change in contact probability between wild type and Δ*hda-1* strains (bottom), between the centromeres and right chromosomal arm, displayed at 10 kb resolution. IGV images of H3K27me2/3 (green) or H3K9me3 (blue) ChIP-seq tracks of wild type and Δ*hda-1* (NKL9) strains are shown below. Genomic distances of LG II (panel **D**) and LG V (panel **H**) are shown above.

### Epigenomic instability in ***Δ***hda-1 leads to progressive decay of histone PTM patterns

The two Δ*hda-1* strain isolates examined by Hi-C produced distinct patterns of aberrant H3K27me3. In the Δ*hda-1* NKL9 isolate, we did not observe aberrant H3K27me2/3 enrichment at the centromere of LGV (**Figure 5H, S6**). In contrast, the Δ*hda-1* NKL10 isolate displayed striking enrichment of H3K27me2/3 at this centromere (**Figure S5H, S6**). These data suggest that redistribution of H3K27me3 may occur in a stochastic or progressive manner, giving rise to unique patterns of aberrant H3K27me3 enrichment in isogenic Δ*hda-1* strains. This could also explain the distinct patterns of H3K27me3 enrichment observed in different HCHC-deficient mutants (**Figure 4)**.

If epigenomic defects observed in Δ*hda-1* occurred slowly over many rounds of mitosis, we reasoned that a newly constructed Δ*hda-1* strain might have a moderated phenotype. To test this, we generated a new Δ*hda-1* strain by split-marker transformation of a wild-type recipient strain. Primary transformants were selected, cultured, and screened by PCR to identify a homokaryotic Δ*hda-1* knockout. This homokaryotic transformant (Δ*hda-1* T0) was then passaged by streaking cells on a plate, isolating a single colony, and then expanding that colony on a slant for a total of three passages (**Figure 6A**). We saved each slant and refer to the sequentially passaged strains as Δ*hda-1* P1, P2, and P3. Cells from each slant were then grown for an additional ∼18 hours in liquid medium, and ChIP-seq was performed using antibodies to H3K27me3, H3K36me3, H3K9me3, and Kac. We estimate that each passage corresponds to ∼25 - 40 rounds of mitosis, based on the number of conidiospores in a typical slant (∼10^8^/slant).

**Figure 6.**
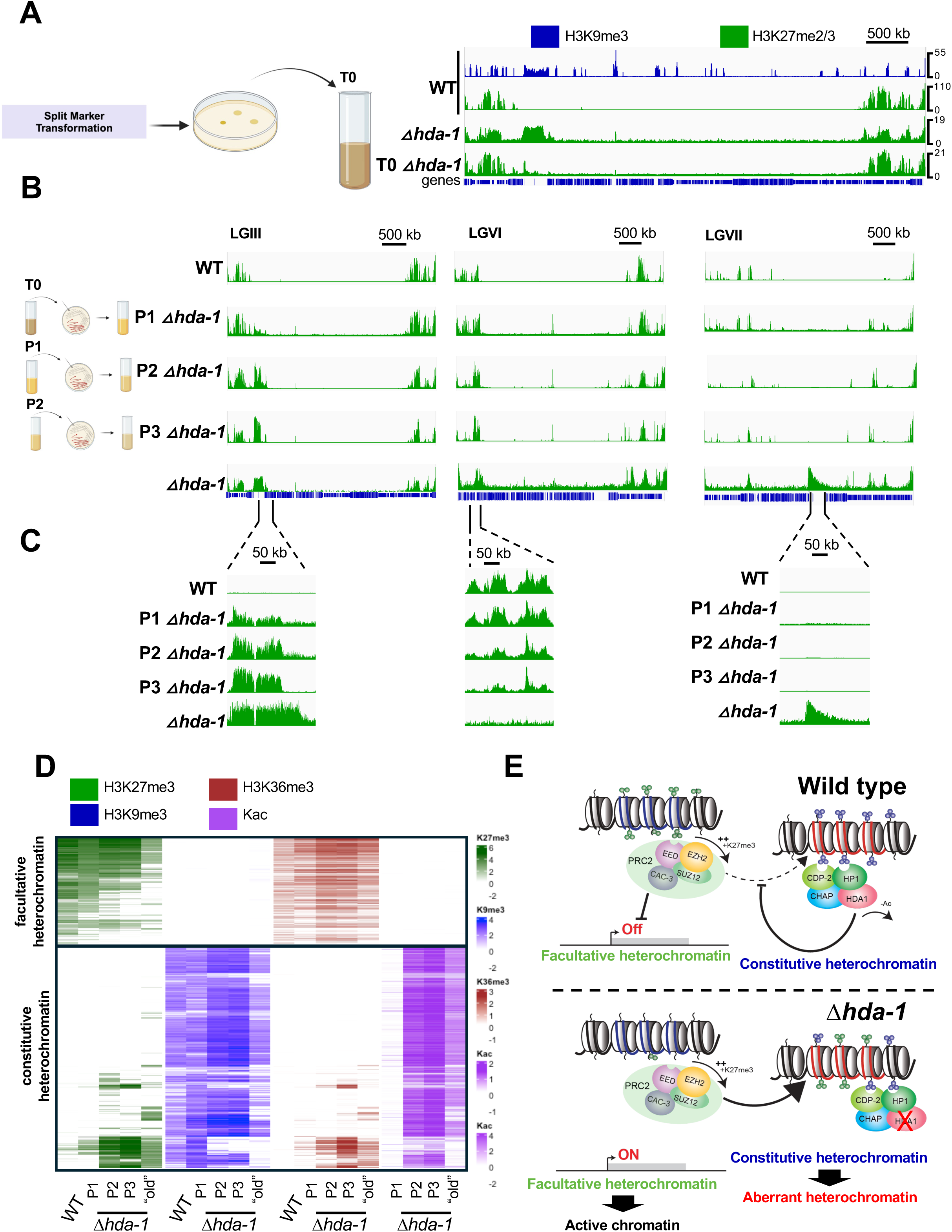
H3K27me3 relocalization is progressive in a laboratory evolution experiment. (**A**) The schematic diagram illustrates the procedure for generating and passaging a new Δ*hda-1* strain (left). The genome browser images (right) show ChIP-seq enrichment of H3K9me3 (blue) or H3K27me3 (green) ChIP-seq enrichment across Linkage Group III for the indicated strains, as in Figure 2A-B. (**B-C**) The genome browser images show H3K27me3 (green) ChIP-seq enrichment across (**B**) Linkage Groups III, VI, and VII for the indicated strains, or (**C**) enhanced images of a cHet region that exhibits progressive gains of H3K27me3 with each passage (left), an fHet region that exhibits progressive loss of H3K27me3 (middle), and the centromere of LGVII (right), which is enriched for H3K27me3 in our original Δ*hda-1* strain but not in the newly constructed and passaged Δ*hda-1* strain. (**D**) The heatmap shows relative enrichment (log2[IP/input]) of H3K27me3 (green), H3K9me3 (blue), or H3K36me3 (orange) in 300bp windows spanning all wildtype fHet and cHet domains for the indicated strains. Relative enrichment of Kac (purple; log2[Δ*hda-1* IP / WT IP]) is shown for the same regions. Enrichment values that were negative or did not meet a statistical threshold (FDR < 0.1) are shaded in white. (**E**) Model of the relationship between fHet (left) and cHet (right) repression in wild type (top) or Δ*hda-1* (bottom) strains.

H3K27me3 enrichment patterns for the primary transformant and the passaged strains are shown in **Figure 6**. The phenotype of the Δ*hda-1* T0 strain closely resembled the wild-type strain with respect to H3K27me3 enrichment patterns (**Figure 6A**). With each passage, we observed progressive loss of H3K27me3 from native fHet domains accompanied by gains of H3K27me3 within typical cHet domains (**Figure 6B-C**). However, even after three passages, some cHet domains that were enriched for H3K27me3 in our original Δ*hda-1* lab strain did not gain H3K27me3 in the new Δ*hda-1* P1 – P3 isolates (**Figure 6B,C**; see CENVII)

We next compared H3K27me3, H3K36me3, H3K9me3, and Kac enrichment patterns in our passaged Δ*hda-1* isolates. Enrichment levels of each modification were quantified in 300 basepair windows covering typical fHet and cHet loci. Enrichment values were then plotted in a heatmap for wild type, the Δ*hda-1* P1, P2, P3 isolates, and our original Δ*hda-1* lab strain (S297; obtained from the Fungal Genetics Stock Center) (**Figure 6D**). After the first passage (P1), both H3K27me3 and H3K36me3 were retained at native fHet domains, but modest gains of H3K27me3 and H3K36me3 were detected in cHet regions. In addition, the Δ*hda-1* P1 isolate displayed increased Kac and relatively normal H3K9me3 enrichment in cHet domains. With each sequential passage, the number of fHet windows with significant enrichment of H3K27me3 and H3K36me3 decreased, accompanied by an increase in the number of cHet windows enriched for both modifications. Progressive gains of H3K27me3/H3K36me3 were correlated with loss of H3K9me3, suggesting that H3K27me3 ultimately leads to displacement of H3K9me3 or that progressive loss of H3K9me3 allows H3K27me3 deposition. As previously observed (**Figure 3**), cHet regions enriched for H3K27me3 in Δ*hda-1* strains showed reduced levels of Kac compared to cHet regions that did not gain H3K27me3, implying that H3K27me2/3 or H3K36me3 directs deacetylation at these regions. We conclude that loss of HDA-1 leads to progressive epigenomic dysfunction.

## Discussion

In this study, we found that HDA-1, a cHet-specific histone deacetylase, is required for repression of PRC2-methylated genes and for maintaining global epigenome stability in *N. crassa*. In HDA-1-deficient cells, cHet domains displayed hyperacetylation, reduced H3K9me3 and abnormal enrichment of two modifications typically enriched in fHet regions, H3K27me2/3 and H3K36me3. Aberrant gains of H3K27me2/3 and H3K36me3 were accompanied by loss of H3K27me2/3 from typical fHet regions and activation of genes normally repressed by PRC2 (**Figure 6E**). These findings extend previous reports of altered H3K27me2/3 in mutants lacking H3K9me3 or HP1, two components required for targeting HDA-1 to cHet domains (35, 36). Notably, our findings differ from those of Jamieson and colleagues, who reported that HCHC components HDA-1 and CDP-2 were not required for normal H3K27me3 patterns (36). This discrepancy likely reflects a difference in the “ages” of Δ*hda-1* and Δ*cdp-2* strains investigated in each study. Here, we showed that redistribution of H3K27me2/3 in a newly constructed Δ*hda-1* mutant occurred gradually, taking place over hundreds of nuclear divisions. It is likely that Jamieson and colleagues performed experiments with Δ*hda-1* and Δ*cdp-2* isolates that were recently backcrossed. If so, relatively normal H3K27me2/3 patterns are expected due the delayed onset of epigenomic defects in the absence of HDA-1 activity. We found that CHAP, an additional HCHC component, was not required for proper H3K27me3. We do not know how many times the Δ*chap* mutant strain obtained from the Neurospora knockout project has been passaged. It is therefore possible that continued propagation of this strain will reveal gradual epigenome instability as observed in Δ*hda-1* and Δ*cdp-2* strains. However, this is unlikely given that previous studies of Δ*chap* strains reported only modest effects on Kac and 3D genome structure compared to other HCHC components, consistent with a non-essential role for CHAP in targeting HDA-1 (42, 56). More generally, our data highlight the importance of controlling for strain age, especially for the study of self-reinforcing repressive chromatin states.

Our results suggest that the aberrant H3K36me3 or H3K27me3 observed at cHet regions of HDA-1-deficient strains directs a compensatory deacetylase activity. In Δ*hda-1* strains, acetylation levels were high in cHet regions that retained normal H3K9me3 enrichment and low in cHet regions enriched for H3K36me3/H3K27me3. This compensatory HDAC activity is likely provided by HDA-3. The *hda-3* gene is essential and therefore was not tested in our screen; however, the HDA-3-containing RPD3L complex was recently identified as a key repressor of PRC2 target genes (80). Thus, we propose that aberrant H3K27me2/3 and/or H3K36me3 targets RPD3L deacetylase activity to a subset of cHet regions in HDA-1-deficient cells. Perhaps depletion of RPD3L from normally repressed fHet is responsible for increased transcription of fHet genes in the Δ*hda-1* mutant. Future experiments are needed to establish how these fHet modifications target the RPDL3 complex.

Our data indicate that HDA-1 normally suppresses off-target activity of PRC2 and ASH-1 at cHet loci. It is possible that HDA-1 functions to restrict DNA accessibility in cHet domains and that loss of HDA-1 leads to transient exposure of cryptic PRC2 or ASH1 recruitment sites. Prior work in *N. crassa* showed that HDA-1 limits DNA methyltransferase accessibility in cHet regions (42). In *S. pombe,* histone deacetylation by the HDA-1 homolog Clr3 is critical for inhibiting chromatin remodeling activity and suppressing nucleosome turnover (88, 89). How PCR2 is recruited to target loci is not well understood in *N. crassa,* but H3K27me2/3 can be directed by telomere repeat sequences in the fungus (90). Future work is needed to determine if the fHet machinery is recruited to cHet regions by DNA sequences or structures that are transiently unmasked in the absence of HDA-1. Alternatively, HDA-1-deficient cells may express cHet-dervied RNAs that recruit PRC2 or ASH1. The ability of PRC2 to bind RNA is conserved from fungi to humans (91), but RNA-dependent recruitment of PRC2 remains controversial (92, 93). A third possibility is that HDA-1 loss may produce changes in the localization or biophysical properties of cHet-associated molecular condensates, leading to off-target activity of PRC2 or ASH1. Phase-separated condensates modulate the activity of PRC2 in *Cryptococcus neoformans.* In *Ccc1*Δ mutants, H3K27me3 accumulates at centromeric cHet domains, where HP1 is proposed to form condensates that concentrate PRC2 to drive off-target methylation in this mutant background (94, 95). In contrast to the situation in *C. neoformans, N. crassa* HP1 acts to prevent aberrant H3K27me3 deposition (35, 36). In Δ*hda-1,* cHet regions that gain H3K27me3 are depleted for H3K9me3 and presumably for HP1. Thus, changes to, or loss of, HP1 condensates may be a precursor to aberrant H3K27me3/H3K36me3 deposition at cHet domains. Additional work is needed to determine how H3K27me3/H3K36me3 are excluded from cHet in wild-type cells and how these PTMs are aberrantly targeted to cHet regions in Δ*hda-1* strains.

Our findings add to the growing body of evidence that HDACs are critical for maintaining epigenome stability. HDACs are required for epigenetic inheritance of H3K9me3 in *S. pombe* (96). It is possible that cHet-specific HDACs maintain proper H3K27me3 patterns in higher eukaryotes, as observed here for *N. crassa*. In mammals, patterns of H3K27me3 are influenced by key features of cHet in the early embryo (97) and by 5mC in a variety of developmental contexts (98). In mESCs, 5mC depletion drives aberrant binding of ncPRC1.1 and deposition of H3K27me3 at pericentromeric heterochromatin and other CpG-rich regions (53, 54). Although DNA methylation is not required for normal H3K27me3 patterns in *N. crassa* (35, 36), antagonism between cHet and H3K27me3 is a shared feature of fungal and mammalian cells. In mESCs, acute 5mC inhibition leads to gradual gains in H3K27me3 at transposable elements and pericentromeric heterochromatin over a two-week period (51). The progressive epigenomic decay observed in 5mC-depleted mESC cells is similar to that observed for HDA-1-deficient cells of *N. crassa*. We speculate that hypoacetylated histones are important for maintaining proper H3K27me3 patterns in both systems. In mammals, it is known that methyl binding domain proteins (MBDs) direct HDAC activity to methylated CpGs (99–101). As such, 5mC depletion in mESCs may lead to hyperacetylation at CpG-dense cHet regions, which could contribute to H3K27me3 redistribution. More generally, data from diverse systems point to HDACs as critical regulators of epigenome stability. Elucidating HDAC-dependent mechanisms that maintain the epigenome may ultimately provide new insights into the origins of epigenetic disorders, including cancer, and could lead to improvements in the detection and treatment of diseases associated with PcG dysfunction.

## Data Availability

All sequencing data are available through the NCBI Gene Expression at the National Center for Biotechnology Information (NCBI) under accession numbers GSE286914 and GSE286535 (ChIP-seq), GSE286916 (CUT&RUN), GSE286538 (Hi-C). Accession numbers for RNA-seq data are included in Supplementary Dataset S2.

## Acknowledgements

The authors would like to Dr. Mary Goll (University of Georgia) and members of the Lewis and Klocko labs for constructive comments on the manuscript. We also thank colleagues at the University of Georgia and the UCCS Department of Chemistry & Biochemistry for helpful discussions.

