## Supplemental Figures and Tables for "Histone deacetylase-1 is required for epigenome stability in *Neurospora crassa*"

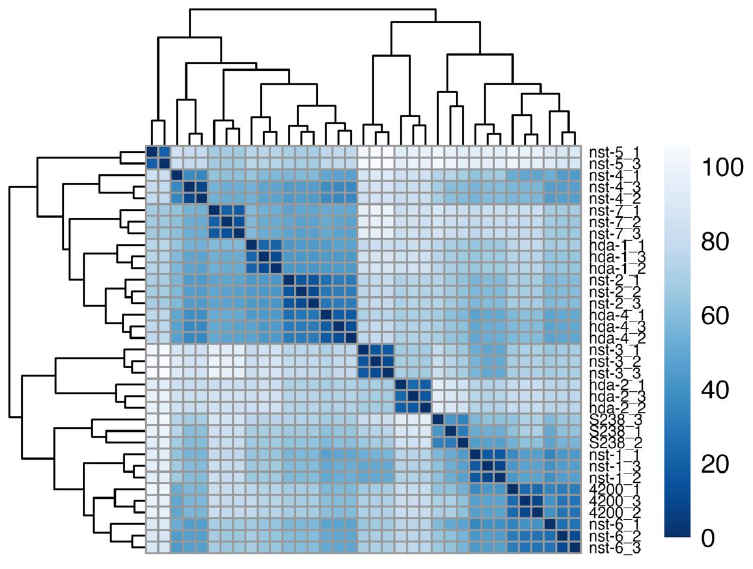

**Figure S1. Replicate similarity for RNA-seq samples analyzed in Figure 1.** Sample distances were determined using DESeq2 and plotted using the R-package pheatmap. Each sample is indicated on the right with the genotype followed by the replicate sample.

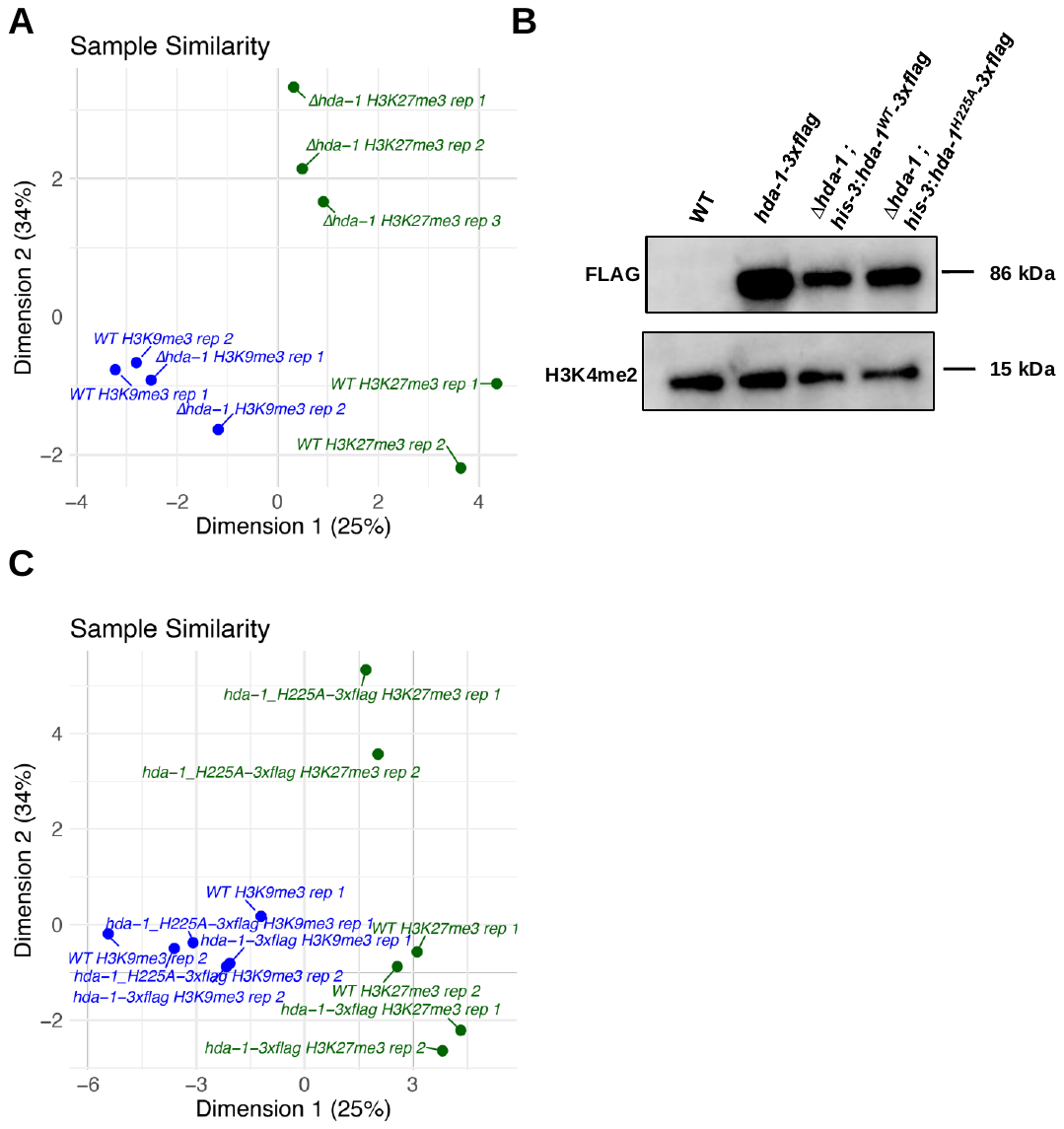

**Figure S2. Replicate similarity and protein expression for samples analyzed in Figure 2.** (**A**) The similarity of replicate samples for wild type and Δ*hda-1* is shown for CUT&RUN experiments performed with antibodies to H3K9me3 (blue) and H3K27me3 (green). Similarity was determined in R using the ‘plotMDS’ function of the csaw package. The genotype and replicate numbers for each point are labeled. (**B**). Western blots of total protein extract from the indicated strains were probed with the indicated antibody. The FLAG signal corresponds to HDA-1-3xFLAG and an antibody to H3K4me2 was used as a loading control. (**C**) The similarity of replicate ChIP-seq experiments using antibodies to H3K9me3 (blue) and H3K27me3 (green) was determined in R using the ‘plotMDS’ function of the csaw package. The genotype and replicate numbers for each point are labeled.

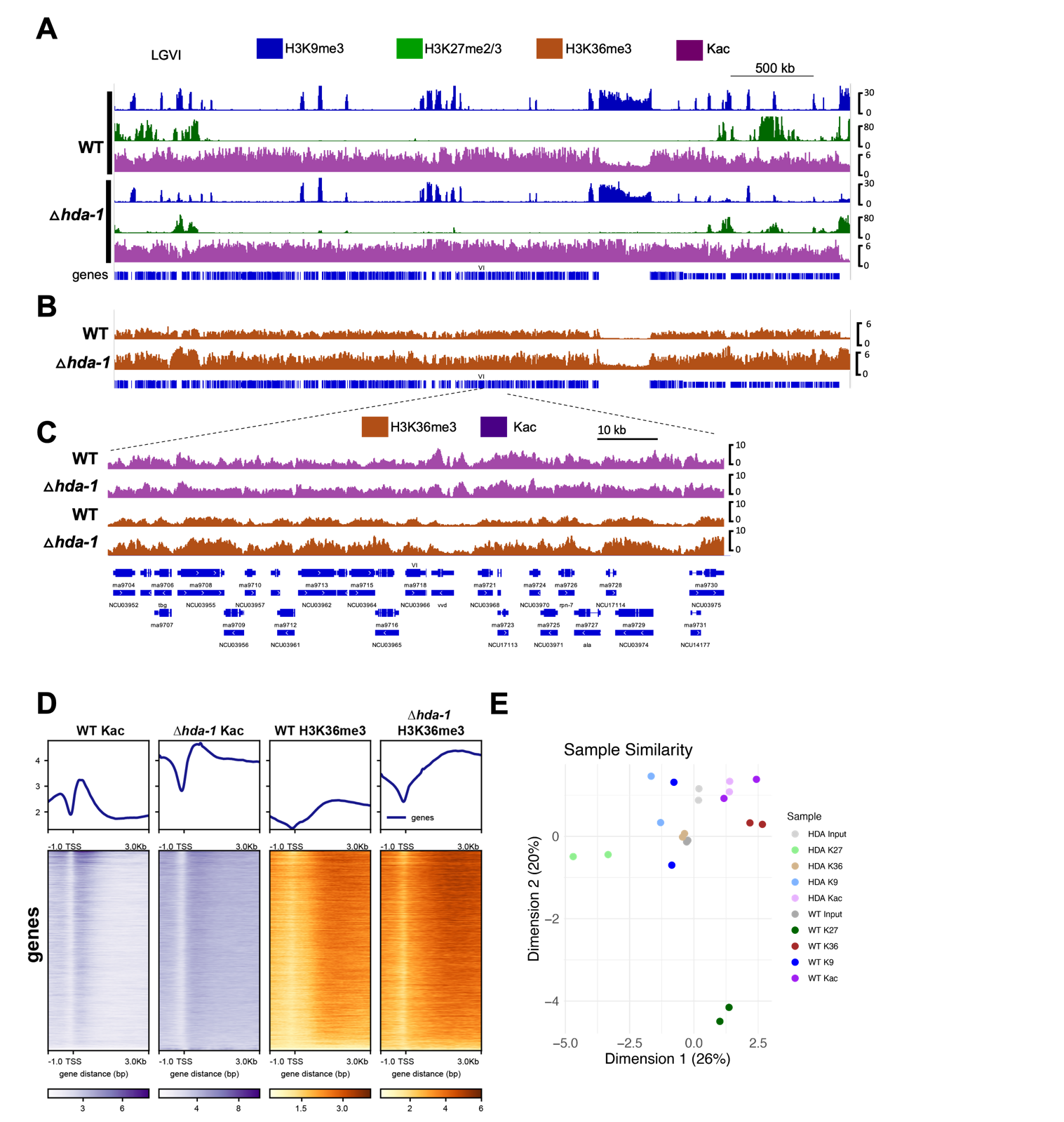

**Figure S3. Replicate data and protein expression for samples analyzed in Figure 2.** (**A**) The genome browser tracks show ChIP-seq enrichment of (**A**) H3K9me3 (blue), H3K27me3 (green), and pan-acetyl lysine (Kac; purple) or (**B**) H3K36me3 across Linkage Group VI. The scale bar indicates genomic distance in kilobase pairs. (**C-D**) The ∆*hda-1* deficient mutant exhibits normal patterns of H3K36me3 and Kac in euchromatin. (**C**) The zoomed-in genome browser image shows ChIP-seq enrichment of Kac and H3K36me3 across a euchromatic region of LGVI in the indicated strains. (**D**) The metaplots and heatmaps show enrichment patterns of Kac (purple) and H3K36me3 (orange) across the transcriptional start sites of all *N. crassa* genes. We observed differences in ChIP efficiency, but similar patterns of enrichment were observed in both strains. **E.** The similarity of replicate samples for ChIP-seq experiments was determined with R using the ‘plotMDS’ function of the csaw package . Two replicates were examined for each condition.

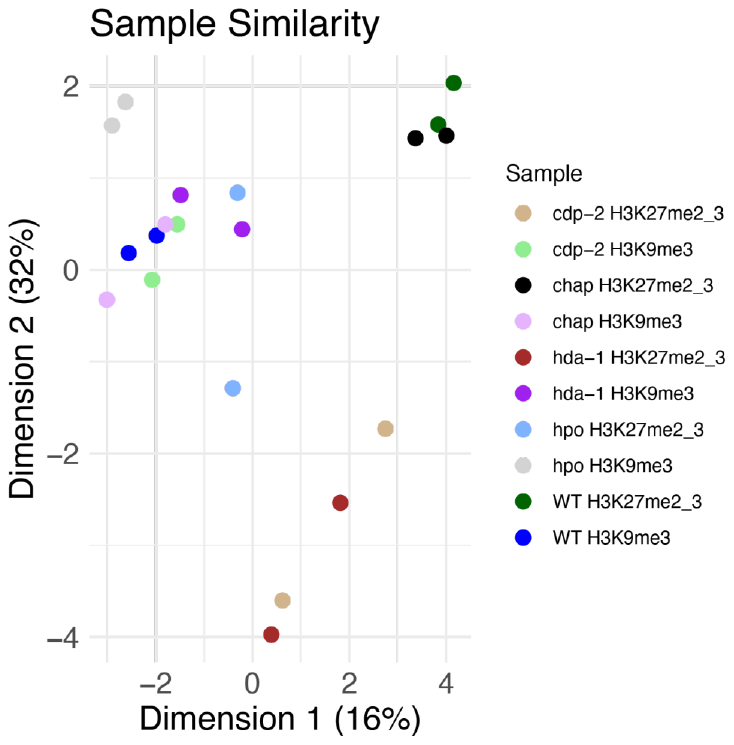

**Figure S4. Replicate CUT&RUN experiments for HCHC-deficient mutants yield reproducible results.** The similarity of replicate samples for CUT&RUN experiments was determined using the ‘plotMDS’ function of the R csaw package. Two replicates were examined for each condition.

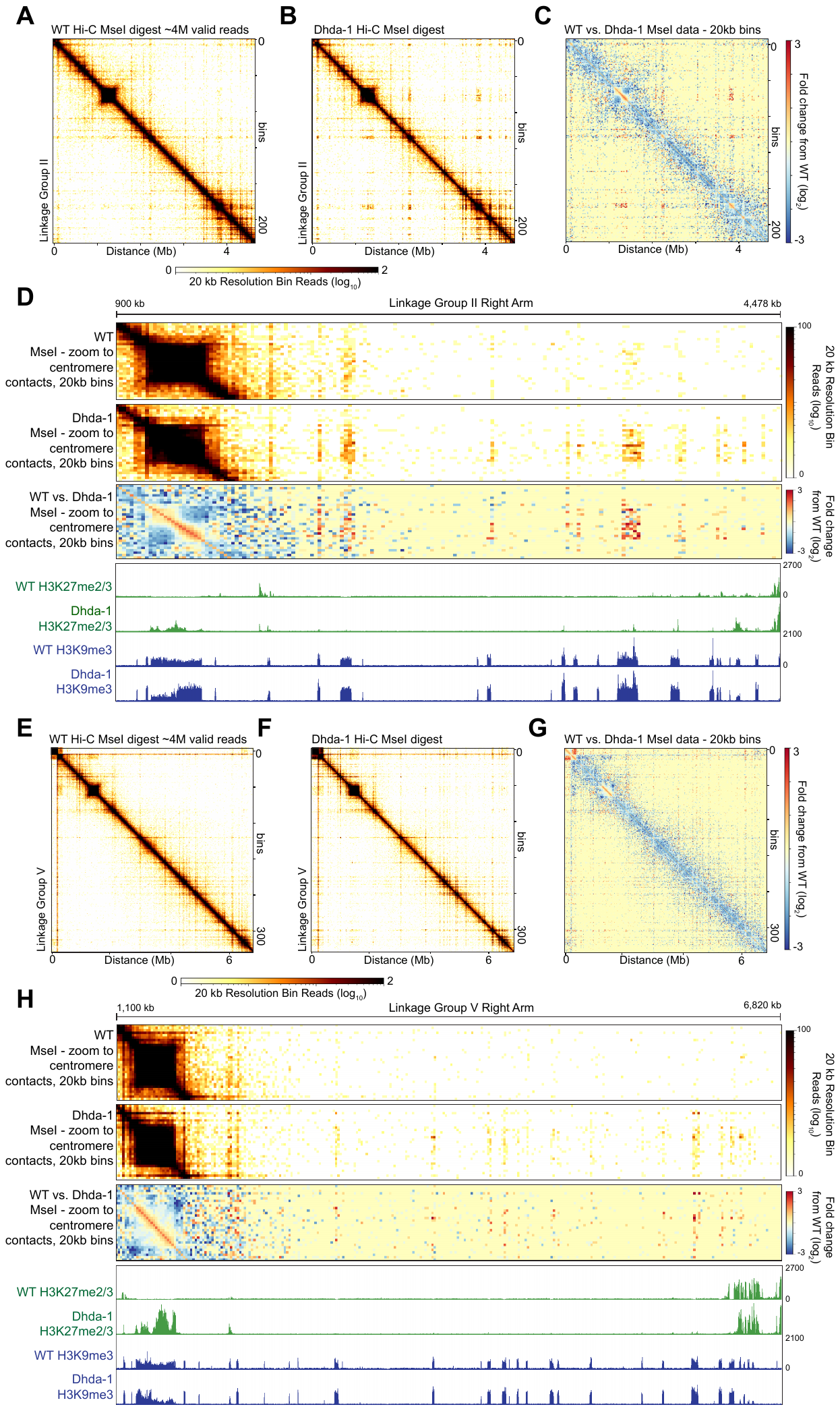

**Figure S5: A replicate Hi-C experiment shows a similar 3D contact profile, but differential H3K27me3 enrichment in a second Δ*hda-1* strain isolate NLK10.** Heterochromatin-specific (*Mse*I) *in situ* Hi-C and H3K27me2/3 or H3K9me3 ChIP-seq of a wild type strain or a ∆*hda-1* strain, NKL9, are displayed across two Linkage Groups (LG), LG II (**A-D**) or LG V (**E-H**). Contact probability heatmaps of raw *in situ* Hi-C reads at 20 kb resolution of a wild-type strain (**A, E**) or the NKL9 ∆*hda-1* strain (**B, F**) are shown; the heatmap scalebar is below, while the genomic distance is indicated on the x-axis and the number of bins indicated on the right. **(C, G**) The comparison heatmaps of the contact probability change (log_2_) between wild type and ∆*hda-1* strains are shown; the heatmap scalebar is to the right. The purple line shows the centromeric region highlighted in panels D and H. (**D, H**) Enhanced contact probability heatmaps of a wild-type strain (top), the ∆*hda-1* NKL9 strain (middle), and the change in contact probability between the wild type and ∆*hda-1* strains (bottom) between the centromeres and right chromosomal arm are displayed at 10 kb resolution. IGV images of H3K27me2/3 (green) and H3K9me3 (blue) ChIP-seq tracks of wild type and ∆*hda-1* strains of the same region are shown below. Distances of LG II (panel **D**) and LG V (panel **H**) are shown above.

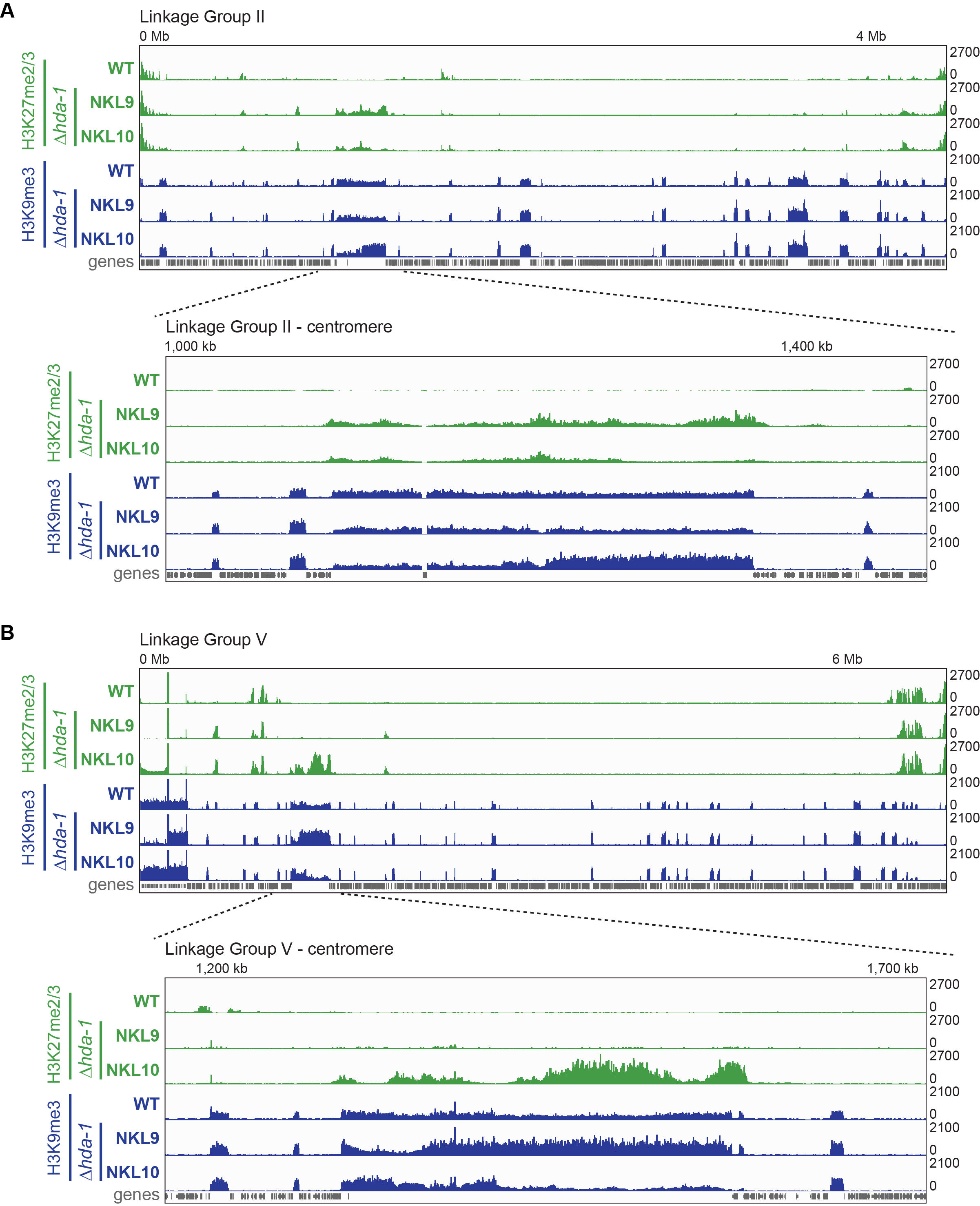

**Figure S6: Variability in H3K27me2/3 enrichment in two ∆*hda-1* progeny. (**A-B) IGV images showing H3K27me2/3 (green) or H3K9me3 (blue) ChIP-seq enrichment across (A) Linkage Group II or (B)Linkage Group V for a wild-type (WT) strain or two∆*hda-1* progeny, NKL9 and NKL10, derived from the same cross, as in Figure 2A. Enhanced images below show the centromeres of each chromosome.

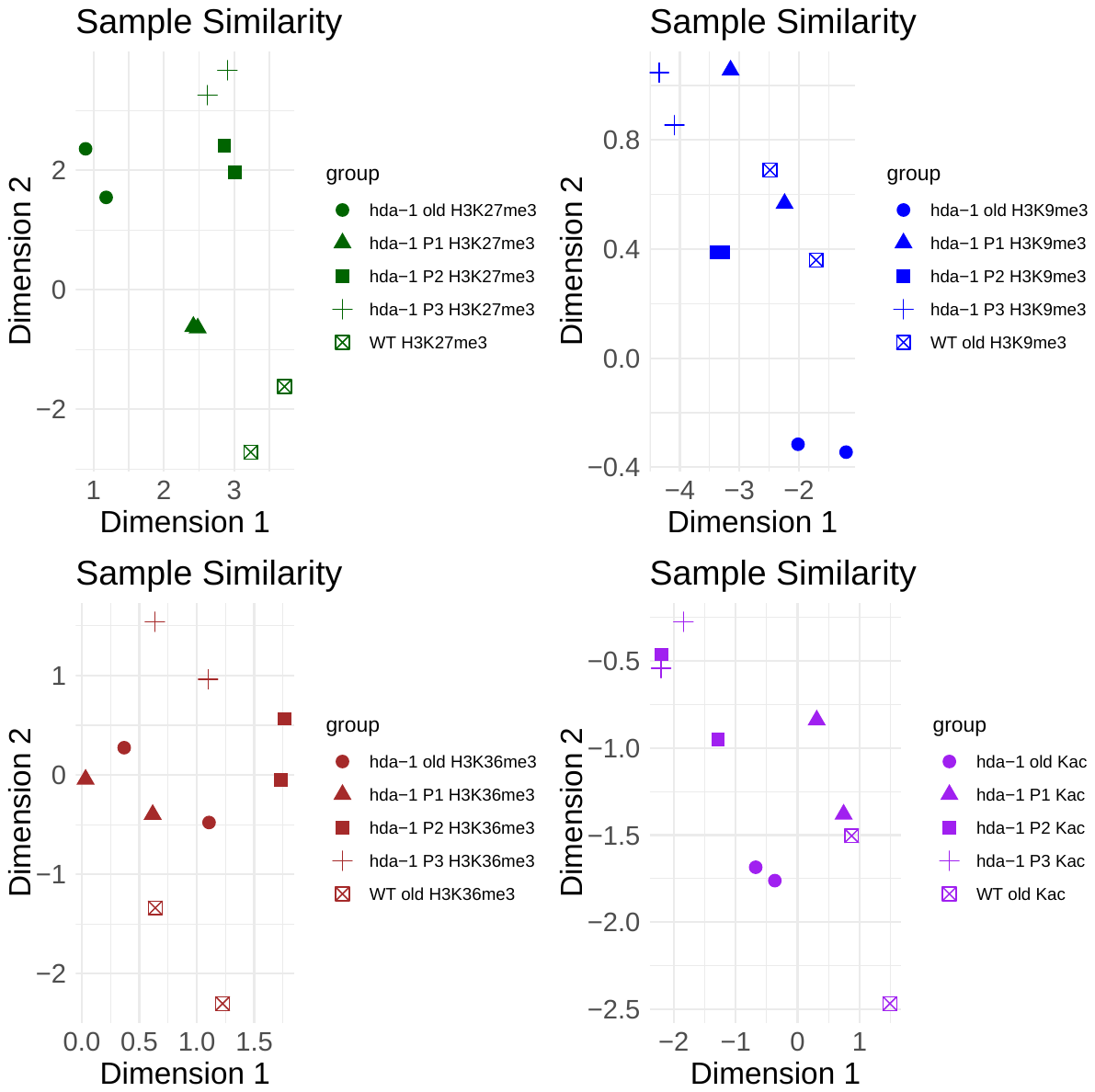

**Figure S7. Replicate ChIP-seq samples for passaged ∆*hda-1* strains and control samples produced similar results.** The similarity of replicate samples for CUTNRUN experiments was determined using the ‘plotMDS’ function of the R csaw package. Two replicates were examined for each strain or condition.

**Table S1. Strains used in this study.**

| Strain | Genotype | Source |
| --- | --- | --- |
| S2 | Wild type *N. crassa* *mat A* (FGSC2489) | FGSC, Kansas City |
| S297 | *hda-1::hph* (FGSC 12003) | This study(1) |
| S181 | *hpo::hph* (FGSC 14522) | (2) |
| S188 | *his-3- hda-1::hph;A* | (3) |
| S189 | *his-3- hda-1::hph;a* | (3) |
| S298 | *cdp-2::hph* (FGSC 11771) | (2)FGSC, Kansas City |
| S299 | *CHAP::hph* (FGSC 12802) | (2) |
| S869 | ∆*hda-1::hph*; *his-3^+^::hda-1^+^::3xFLAG* | This study |
| S870 | ∆*hda-1::hph*; *his-3^+^::hda-1^H225A^::3xFLAG* | This study |
| NKL9 | Δ*hda-1* (backcross isolate) | This study |
| NKL10 | ∆*hda-1* (backcross iolate | This study |
| S871 | ∆*hda-1::hph* (newly generated) | This study |
| S875 | S871 passage 1 | This study |
| S976 | S871 passage 2 | This study |
| S877 | S871 passage 3 | This study |

**Table S2. Primers used in this study**

| **Name** | **Sequence** |
| --- | --- |
| HDA-1 5’ Flank FP | CGATCCGTCCCCCCCAA |
| HDA-1 3’ Flank FP | TTCGAGGCGGTTGTTGGTGA |
| SM1 | AAAAAGCCTGAACTCACCGCGACG |
| SM2 | TCGCCTCGCTCCAGTCAATGACC |
| pBM61_Inverse P3 | ATACGACTCACTATAGGGCGAATTGGAGCT |
| pBM61_Inverse P2 | GTGGACGGCTAATGGGGTCTGAATGCTAAA |

**Table S3. Antibodies Used in this study**

| **Antigen** | **Supplier** | **Catalog #** |
| --- | --- | --- |
| H3K27me2/3 | Active Motif | 39535 |
| H3K9me3 | Active Motif | 39162 |
| H3K36me3 | Abcam | ab9050 |
| Pan-acetyl-lysine | Cell Signaling Technology | 9441S |
| FLAG | Sigma-Aldrich | F1804 |
